# NMR backbone assignment and dynamics of Profilin from Heimdallarchaeota

**DOI:** 10.1101/2020.04.20.050468

**Authors:** Syed Razaul Haq, Sabeen Survery, Fredrik Hurtig, Ann-Christin Lindås, Celestine N. Chi

## Abstract

The origin of the eukaryotic cell is an unsettled scientific question. The Asgard superphylum has emerged as a compelling target for studying eukaryogenesis due to the previously unseen diversity of eukaryotic signature proteins. However, our knowledge about these proteins is still relegated to metagenomic data and very little is known about their structural properties. Additionally, it is still unclear if these proteins are functionally homologous to their eukaryotic counterparts. Here, we expressed, purified and structurally characterized profilin from Heimdallarchaeota in the Asgard superphylum. The structural analysis shows that while this profilin possess similar secondary structural elements as eukaryotic profilin, it contains additional secondary structural elements that could be critical for its function and an indication of divergent evolution.

## Biological context

The origin of the eukaryotic cell remains an unsettled scientific question and several hypotheses have been put forward to explain the complex evolutionary history of the eukaryotic cell^1 2 3 4^. The Woese hypothesis proposes three domains of life - Archaea, Bacteria and Eukaryota, with independent evolutionary trajectories^5^. The eocyte hypothesis suggests the existence of only two domains - bacteria and archaea, and that eukaryotes emerged from the symbiotic relationship of an unknown archaeal host with an alphaproteobacterium^6^. Recently, environmental metagenomic sampling led to the discovery of the Asgard superphylum. Comparative genomic analysis of Asgard archaea and eukaryotes appears to support the eocyte hypothesis^7^. The genomes of Asgardarchaea are enriched with proteins previously considered eukaryote-specific, so called eukaryotic signature proteins (ESPs), and phylogenetic analysis placed the Asgardarchaea in a monophyletic group with eukaryotes^8^.

Actin plays a crucial part of the eukaryotic cytoskeleton and is essential to many processes, including cellular motility, cell division, endocytosis, intracellular cargo transport, amongst many other^9^. Because of the central role actin plays in the eukaryotic cell, the sequence of actin remains highly conserved among eukaryotes. While actin homologues are widespread throughout all domains of life, the dynamic actin cytoskeleton and the regulatory actin-binding proteins are a hallmark of eukaryotic life.

The Asgard genomes contain close actin homologues and several actin-binding proteins; including profilin, gelsolin, Arp2/3 complex subunit 4 and a large family of small GTPases that regulate the actin cytoskeleton in eukarotes^8^. This posits the question; do these archaea possess an actin cytoskeleton with complex regulation analogous with the eukaryotic cytoskeleton? While metagenomic analysis has identified these proteins, their cellular function is still poorly understood. Laboratory culturing of these organisms is still in early development which makes *in vivo* comparison with eukaryotic homologs difficult^10^. Currently, protein production in heterologous expression systems and reconstitution of the purified complexes *in vitro* represents one of the best approaches in characterizing their function. Profilin is expressed in most, if not all, eukaryotic cells and is one of the most important proteins in regulating actin cytoskeletal dynamics^11^. Eukaryotic profilin (eprofilin) is a small protein (approximately 14-19 kDa) which sequester monomeric G-actin from the cytoplasmic pool, thus controlling polymerization^12^. Despite significant divergence at the sequence level, the eprofilin tertiary structure is well-conserved and folds into 3D structures constituting 7 β-strands and 4 α-helices^13^. eprofilin promotes the elongation of actin filament assembly at the barbed end by acting as a nucleotide exchange factor, and by interacting with elongation factors such as Ena/Vasp, Formins, and Wasp^14–16^. These nucleation factors bind eprofilin through a polyproline motif at a domain physically separate from the actin binding-site. Moreover, eprofilin can also bind to phosphatidylinositol 4,5-bisphosphate (PIP_2_)^17^ at the plasma membrane which results in a reduced affinity towards polyproline and actin^17^. eprofilin also competes with phospholipase C for PIP_2_ binding which leads to interference with the PI3K/AKT signaling pathwa^18^.

Recently, it has been shown that profilins encoded in several lineages of the Asgardarchaea not only share structural similarity with eukaryotic orthologues but are able to regulate the function of eukaryotic actin. This implies that profilin from Asgardarchaea have the potential of complex regulation of the hypothetical actin cytoskeleton as well^19^. In contrast to human profilin I, a previous study showed that the Asgard profilins (Loki 1 and 2, Thor, Odin and Heimdall) did not show polyproline binding. This led the authors to suggest that Asgard profilins do not bind polyproline, and that polyproline directed actin assembly is a later addition in eukaryotic evolution^19^. However, PIP_2_ was shown to modulate the affinity of Asgard profilin towards rabbit actin in a functional assay^19^. Nevertheless, some of the Asgard genomes are incomplete and the structural and functional relationships of representative profilins from different Asgard lineages are still poorly understood. It might therefore be too early to assume that Asgard profilins do not bind polyproline. In addition, the crystal structures of various profilins combined with functional data do not only reveal structural similarity between Asgard profilins, but also highlights some subtle differences at the species level^19^. Within the Asgard superphylum, the Heimdallarchaeota appears to currently be the closest relative of eukaryotes^7^. Here we present the NMR backbone assignment and dynamics of the Heimdallarchaeota profilin (heimProfilin) as a first step towards characterizing it structurally. These NMR amino acid specific assignments and dynamics provide for the first time an atomic snapshot of heimProfilin as well as providing further evidence for the idea that the Asgard encoded proteins possess similar structural elements and are likely to perform similar roles as those in eukaryotes.

## Methods and experiments

### Protein expression and purification

Heimdallarchaeota profilin (GenBank: OLS22855.1) was cloned into the pSUMO-YHRC vector, kindly provided by Claes Andréasson (Addgene Plasmid #54336; RRID: Addgene_54336) with an N-terminal 6xHistidine-tag and a SUMO-tag (cleavable with Ulp1 protease). The vector was transformed and expressed in *E. coli* Rosetta DE3 cells. Initially, the cells were grown in 2x TY media at 37 °C until the optical density of the culture was 0.8 at 600 nm. Cells were then harvested by centrifugation at 4000 x g for 15 minutes at 4 °C and washed twice with M9 medium.

The cells were then transferred into M9 media supplemented with 1g/L ^15^N-ammonium chloride and 1g/L ^13^C-glucose and grown for 1 hour at 30 °C. Protein expression was induced by 0.5 mM IPTG. For Deuterium (^2^H) labelling, the M9 medium was prepared with 100% or 50% D_2_O and cells were grown overnight at 30 °C. Post-induction, the cells were harvested by centrifugation and resuspended in the binding buffer (50 mM Tris-HCl pH 7.5, 0.3 M NaCl, 1 mM TCEP, 10 mM imidazole, 10% glycerol). The cells were then lysed by sonication and the cell lysate was clarified by centrifugation at 25,000 x g for 45 min at 4 °C and finally filtered through a 0.2 µM syringe filter (Sarstedt). The supernatant was loaded onto a His GraviTrap column (1ml, GE healthcare) and the bound protein was eluted with binding buffer containing 250 mM imidazole. The protein was incubated with Ulp1 protease overnight at 4 °C to cleave the SUMO-tag including the Histidine-tag. The protein was desalted using a PD10 column (GE Healthcare) and loaded onto a His GraviTrap column again to remove the tag and the Ulp1 protease. The protein was concentrated using a 10,000 NMWL cutoff centrifugal filter (Merck Millipore) and further purified on a Superdex 75 10/300 GL (GE Healthcare) size exclusion column, equilibrated with 25 mM Tris-HCl, 50 mM NaCl, 5% Glycerol, 1 mM TCEP at pH 7.5. Protein concentration was determined using the molar absorption coefficient at 280 nm (29450/M/cm).

### NMR Spectroscopy

Double labeled ^15^N, ^13^C, or triple labeled ^15^N, ^13^C, ^2^H were prepared to a concentration of 20 mg/ml in 25 mM Tris-HCl, 50 mM NaCl, 5% Glycerol, 1 mM TCEP at pH 7.5 and thereafter supplemented with 3% D_2_O and 0.03% sodium azide. The NMR assignment experiments were performed at 308K on a triple-resonance Bruker 900, 700 or 600 MHz spectrometers equipped with a cryogenic probe. NMR relaxation experiments were performed on a 600 MHz spectrometer at 298 K. Backbone sequence-specific assignments were carried out using the following experiments 2D ^1^H-^15^N-TROSY, 3D TROSY-HNCACCB, 3D TROSY-HNCA, 3D TROSY-CO) CACB and 3D TROSY-HN(CO)CA. For side-chain assignments, 2D ^1^H-^13^C CT-HSQC, 3D HBHA(CO)NH and 3D HCCH-TOCSY spectra were utilized. For assignment and fold verification 3D NOESY as well as ^3^*J*_*HNHα*_ for secondary structure verification were measured. For Backbone *R*_1_, *R*_2_ rates and hetero-nuclear NOES were determined in an interleaved manner with the experiments from the Bruker pulse program library. For *R*_1_ and *R*_2_ rates, the relaxation delay was sampled for 9 and 8 delay-durations which were pseudo-randomized, respectively (*R*_1_: 20, 60, 100, 200, 400, 600, 800 and 1000 ms and *R*_2_: 16, 33, 67, 136, 170, 203, 237 and 271 ms). The relaxation delay time was up to 1.5 s for *R*_1_ and 1 s for *R*_2_. The [^1^H]^15^N-hetNOE experiment and a reference spectra were recorded with a total 2 s ^1^H saturation time for the NOE experiment and the same recovery time for the reference experiment. The order parameter S^2^ and the internal correlation time were calculated with the program dynamic center. The rotational diffusion tensor was estimated from the ratio of the relaxation rates (R and R). TALOS and CYANA were employed to predict secondary structure, using ^1^H^N^, ^15^N, and ^13^C^α^ chemical shifts. All other data were processed with topspin and analyzed using CCPNMR^20^ and CYANA^21^.

### Assignment and data deposition

The expressed and purified heimProfilin corresponds to the full length as was generated from metagenomics data^8^. It consists of 148 amino acids which was purified with a cleavable tag that leaves no additional N-terminal amino acids (see methods). This profilin possesses a 20-amino acid extension compared with the previously characterized eprofilins or those from Loki I and II and Odin. We obtained up to 88% of all backbone and up to 80% of all side-chain assignments. 135 of the 148 non-proline amide residues were assigned in the ^1^H-^15^N TROSY (Fig. 1). The following amides were not possible to assign: M1, K2, D3, I6, K11, K14, I19, S25, E27, N62, S85 and N89. The missing amides could be due to motional broadening or fast solvent exchange. We obtained 92% of the C_α_ and C_β_ resonance assignments. H_β_ and H_α_ proton shifts were completed to 97% and 96%, respectively. These assignments were further verified by ^15^N/^13^C 3D NOESY spectra. Backbone and side-chain chemical shifts assignments have been deposited to the Biological Magnetic Resonance Data Bank (BMRB) with the Accession Number 50190.

**Figure 1:**
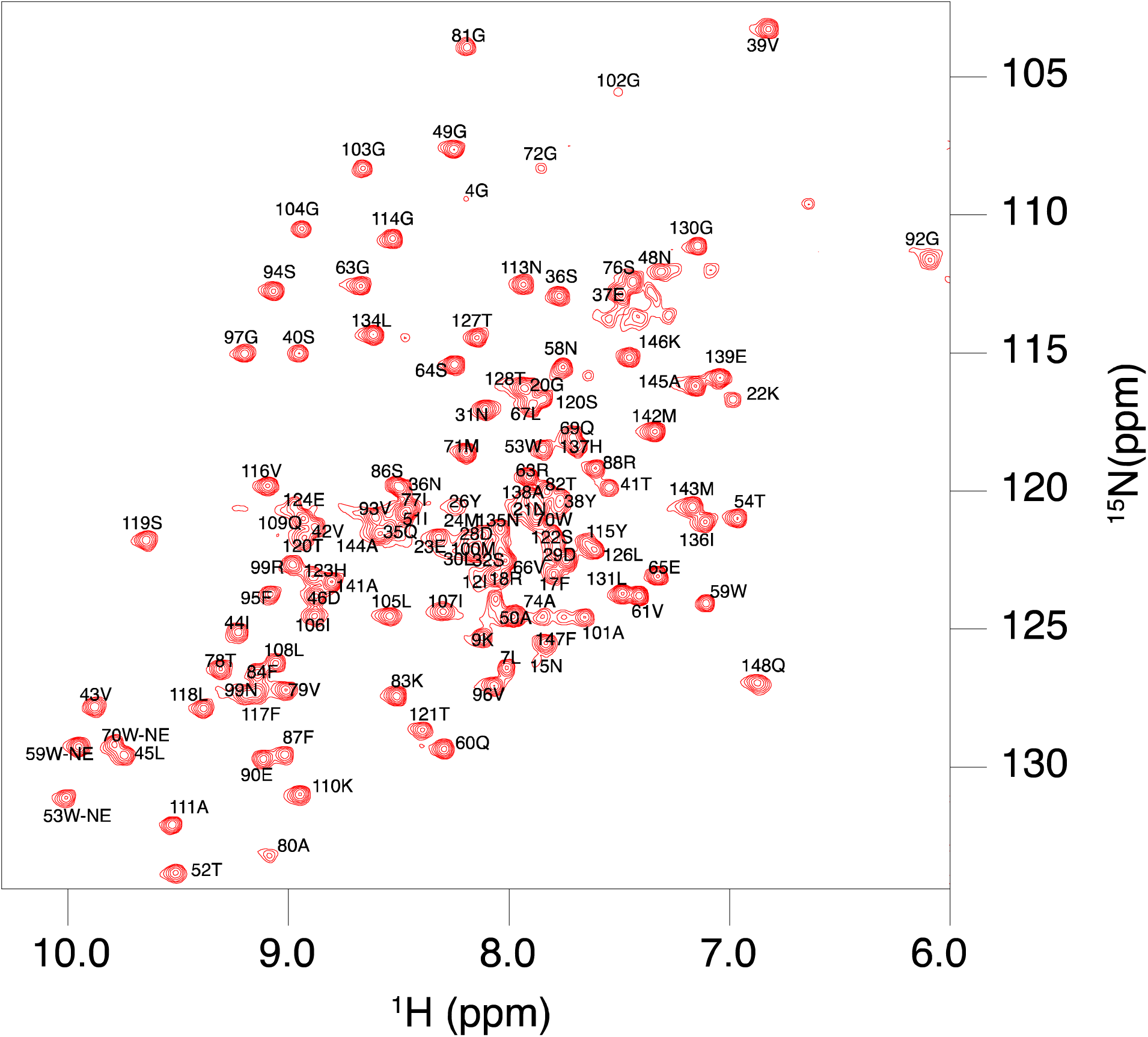
1H-15N TROSY correlation spectrum of Heimdallarchaeota profilin. All ^1^H-^15^N pairs that were assigned in this study. Side-chains of Glutamine and Asparagine are not assigned or shown.

### Secondary structure analysisthe reference experiment

The structures of eukaryotic and Asgard profilin from Loki (1 and 2) and Odin have been determined by X-ray crystallography^19^. However, no structural information is available from the heimProfilin which appears to be the closest relative to the eukaryotes. With the completed assignments, it was now possible to analyze the secondary structure characteristics of this profilin to see if it adopts similar secondary structural elements. Analysis of sequential and medium range NOEs revealed stretches of dNN, dNN(i, i+2), dαβ(i, i+3), dαN(i, i+3). Residues 29– 33, 64-68, 124-127 and 125-144 continual revealed dαN (i, i+4) NOEs, indicating the presence of helices in this region. This is supported by the ^3^*J*_*HNHα*_ coupling constants for these residues which display small values typical of alpha helices (Fig. 2). ^13^C_α_ and C_β_ shifts are frequently used to predict secondary structure propensities. C_α_ shifts generally tend to shift upfield in a beta-sheet and extended strands relative to the random coil values. In alpha helices, these C shifts tend to shift downfield22. For C_β_ values the opposite is true, they shift downfield for beta-sheets and extended strands and upfield for alpha helices. The C_α_ and C_β_ values relative to random coil values are shown in figure 2. Examination of these plots indicates clear helical regions covering residues 29-34, 64-68, 124-127 and 135-144. The helical region between residues 64-68 has not been observed in previous profilin structures. The region of beta strands also agrees with NOEs values and slightly increased 3J values. This analysis indicates that the overall secondary structural elements are preserved from archaea to eukaryotes albeit with some slight differences in their lengths. In addition, we observed an additional helix between residues 64-68 which was not present in the previously determined profilin structures. This might be important for modulating profilin-actin interaction and other physiological roles.

**Figure 2:**
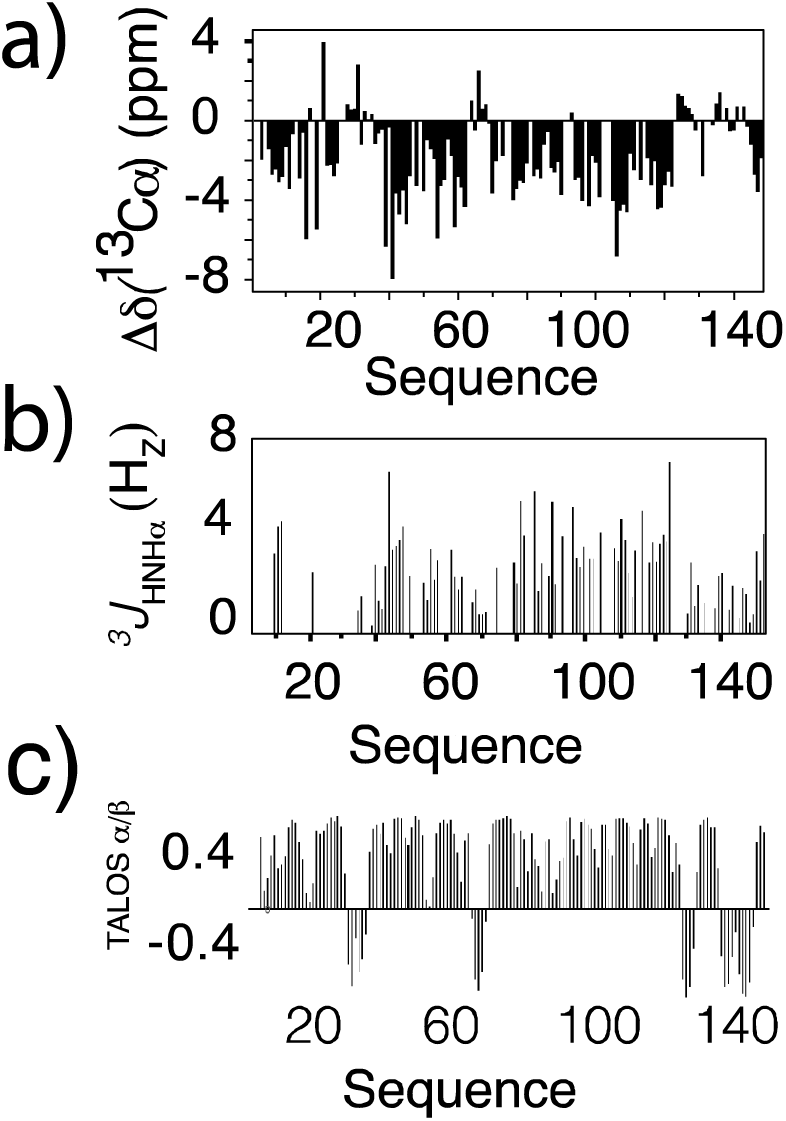
Secondary structure characterization of Heimdallarchaeota profilin. (a) Sequence-specific 13C_α_ secondary chemical shifts (δΔ^13^C_α_) along the amino acid sequence of heimProfilin. (b) ^3^*J*_HNHα_ couplings plotted as a function of amino acid sequence. (c) TALOS secondary structure prediction based on ^1^H, ^15^N and ^13^C_α_ shifts plotted as function of amino acid sequence. All three suggest the presence of helical and extended strands in similar regions. The presence of helices between residues 29-34, 64-68, 124-127 and 135-144 are clearly visible. 3JHNHα couplings are generally lower for helices (2-4 Hz) and higher for beta strands and extended regions (2-8 Hz). Very few ^3^*J*_HNHα_ couplings were obtained for residues 1-20. However, the ^13^C_α_ shifts and the TALOS prediction clearly shows that this region is extended.

### Backbone dynamics

*R*_1_ and *R*_2_ rates in addition to [^1^H]-^15^N hetNOE are frequently used to estimate the flexibility of proteins^23^. Deviation of *R*^1^ and *R*_2_ rates for [^1^H]-^15^N moieties from the average value often indicate a change in motional property. R_1_ values that are larger than the average indicates the presence of flexibility in the ps-ns time range. On the other hand, *R*_2_ rates with higher values than the average indicates regions of slow conformational exchange in the µs-ms time scale. [^1^H]-^15^N hetNOE with negative or near zero values indicate regions of high flexibility with motions faster than approximately 1 ns. We measured and plotted the longitudinal R and transverse R rates as well the [^1^H]-^15^N hetNOE versus amino acids sequence (Fig. 3). Overall, the results from these values indicate a highly rigid protein between residues 25-148 (Fig. 2). However, N-terminal residues 1-24 show a high degree of flexibility, which is reflected in the very low [^1^H]-^15^N hetNOE values (Fig. 2). We also back calculate order parameter S^2^ and internal correlation time τe. A plot of the calculated order parameter *S*^2^ and internal correlation time τe is shown in figure 3d. As shown in the plot, only the N-terminal 1-24 amino acids show some degree of flexibility with very low order parameter and high degree of internal motion. A few residues along the protein sequence indicate some degree of flexibility. We determined the correlation time τc of 11.3 ns. This value is slightly higher for a protein of this size indicating probably due to the extended N-terminal loop not completely structured.

**Figure 3:**
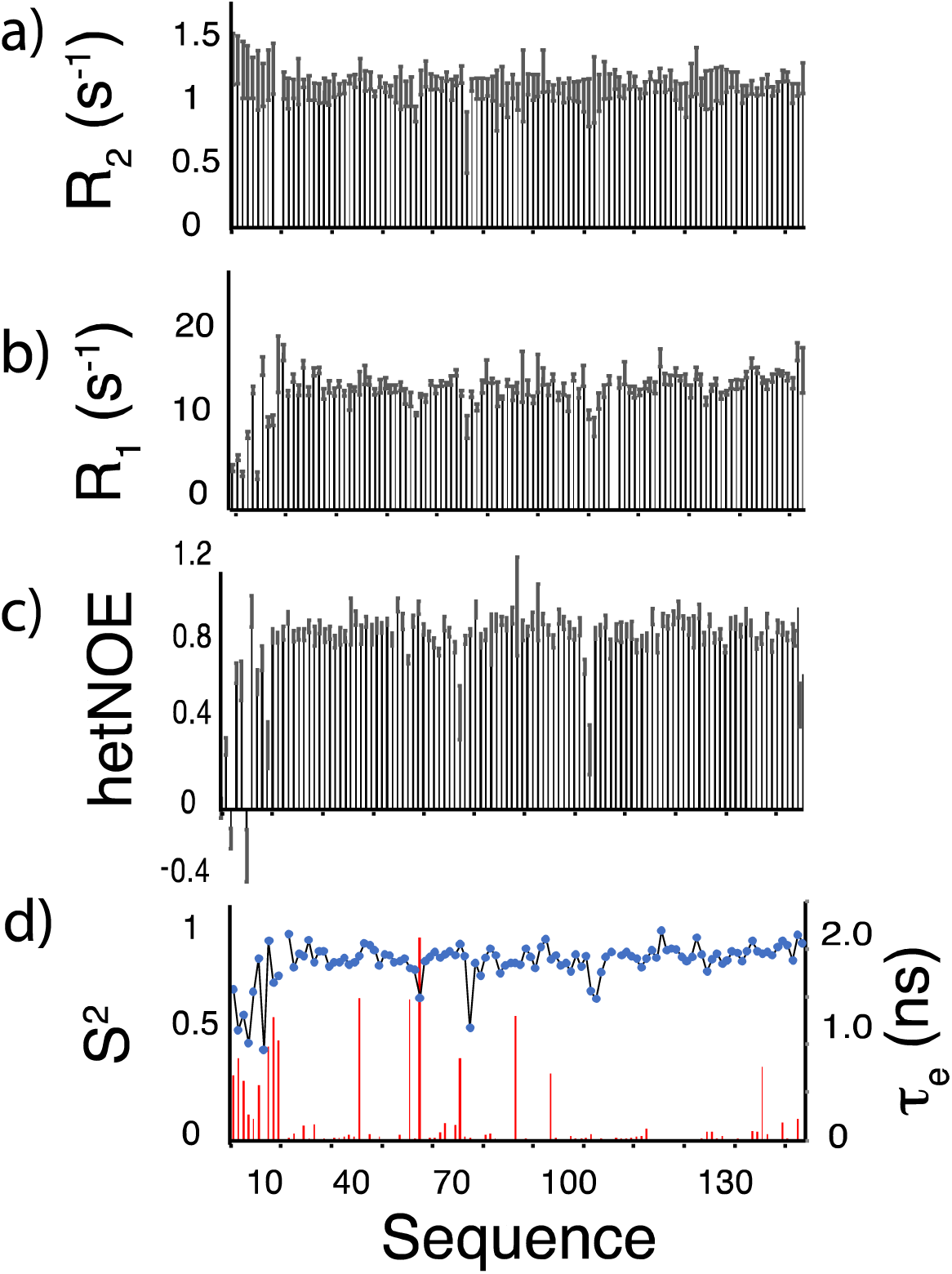
Dynamic characterization of the backbone based on [^1^H-]^15^N. (a) Longitudinal ^15^N *R*_2_ relaxation rates plotted as a function of amino acid sequence. (b) Transverse 15N R relaxation rates versus the amino acid sequence. (c) [^1^H]^15^N-hetero-nuclear NOE data (hetNOE) along the amino acid sequence. (d) calculated *S*^2^ order parameter (left axis) and internal motion τe (right axis) plotted as a function of the amino acid sequence. All parameters indicate a very rigid molecule structure from the N-terminal 20 amino acids which show some degree of ps-ns motion based on the elevated *R*_1_, lower hetNOE and lower *S*^2^ order parameter.

### Conclusions

In this study, we have determined the NMR backbone and dynamic data of a profilin from Heimdallarchaeota in the Asgard superphylum. Our secondary structure analysis indicates that this profilin possess similar structural elements to eukaryotic homologues, all beit at varied lengths. Our data also indicates that the heimProfilin appears rigid apart from N-terminal residues 1-24 which are not present in previously characterized eukaryotic profilins. We observed an additional helix between residue 64-68 which lies in the interface of the actin binding site when compared to eukaryotic profilin, and likely plays a role in modulating acting polymerization.

## Declarations

### Ethics approval

not applicable.

### Consent to participate

not applicable.

### Consent for publication

all authors read an approved the manuscript.

### Availability of data and material

all data and material are available and can be obtain from the authors

